# Towards bone-remodeling-on-a-chip: self-assembling 3D osteoblast-osteoclast coculture in a microfluidic chip

**DOI:** 10.1101/2023.03.11.532167

**Authors:** M.A.M. Vis, F. Zhao, E.S.R. Bodelier, C.M. Bood, J. Bulsink, M. van Doeselaar, H. Eslami Amirabadi, K. Ito, S. Hofmann

## Abstract

Healthy bone is maintained by the process of bone remodeling. An unbalance in this process can lead to pathologies such as osteoporosis which are often studied with animal models. However, data from animals have limited power in predicting the results that will be obtained in human clinical trials. In search for alternatives to animal models, human *in vitro* models are emerging as they address the principle of reduction, refinement, and replacement of animal experiments (3Rs). At the moment, no complete *in vitro* model for bone-remodeling exists. Microfluidic chips offer great possibilities, particularly because of the dynamic culture options, which are crucial for *in vitro* bone formation. In this study, a scaffold free, fully human, 3D microfluidic coculture model of bone remodeling is presented. A bone-on-a-chip coculture system was developed in which human mesenchymal stromal cells differentiated into osteoblasts and self-assembled into scaffold free bone-like tissues with the shape and dimensions of human trabeculae. Human monocytes were able to attach to these tissues and to fuse into multinucleated osteoclast-like cells, establishing the coculture. Furthermore, a set-up was developed allowing for long-term (35 days) on-chip cell culture with benefits including continuous fluid-flow, low bubble formation risk, easy culture medium exchange inside the incubator and live cell imaging options. This on-chip coculture is a crucial advance towards developing *in vitro* bone remodeling models to facilitate drug testing.

## 1. Introduction

Bone is a dynamic tissue with multiple functions, i.e. allowing for movement, protecting organs, and storing essential minerals. Healthy bone is constantly maintained by the process of bone remodeling in which bone tissue is resorbed by osteoclasts and formed by osteoblasts. This process is regulated by osteocytes. Under homeostatic circumstances, the amount of bone matrix that is resorbed is equal to the amount that is formed, and thus bone mass is maintained [1].

In pathologies such as osteoporosis, the balance between formation and resorption is disturbed, leading to changes in the mechanical properties of the bone and their risk for failure. To study bone pathologies, animal models are often used. However, animal models are costly, time-consuming and ethically undesirable [2]. Furthermore, data from animals frequently fail to predict the results obtained in human clinical trials [2], [3]. In search for alternatives to animal models, human *in vitro* models are emerging as they address the principle of reduction, refinement, and replacement of animal experiments (3Rs) [4].

Organ-on-a-chip devices represent one of the recent successes in the search for *in vitro* human models that can recapitulate organ-level and even organism-level functions [2]. These organ-on-a-chips promise several advantages over traditional techniques such as integration of structural and dynamic cues, small amount of cells, samples and reagents leading to decreased costs and options for parallel and real-time analysis. Many successful studies have been published, showing the potential of organ-on-a-chip technologies [5], [6]. However, there are still challenges that need to be overcome.

One of the most important challenges is the formation of air bubbles within the device [7]. Air bubbles can get trapped in the chip and thereby disturb its performance. When air bubbles flow over cells in culture, cell membranes damage can be caused to the due to dynamic air-liquid interfaces. This can result in cell death when the surface tension of these interfaces is high enough to rupture the cell membrane [7]. In addition, bubbles can cause blockage of culture medium perfusion. This blockage could lead to pressure build-up that disturbs the stability of the system, can cause device failure and sudden mechanical stimulation of the cells [8].

Next to the bubble formation, the practical handling of the system may be challenging. Microfluidic technology often promises scale-up possibilities, but in practice this can be difficult to achieve [9], [10]. Setting up the cell culture, connecting the tubing, pumps and chips is usually a manual and complex task, limiting the number of samples that can be processed at the same time. In addition, researchers in the field advocate standardization [10], [11]. It is important that the design of the systems is intuitive and straightforward, while still enabling reliable and robust operation.

In this study, both bubble formation and ease-of-use were tackled by designing a practical experimental set-up for long-term (35 days) on-chip culture. The design requirements were: easy transportation between safety cabinet and incubator, short tubing length between medium reservoir and chip to reduce bubble formation risk, addition of live cell imaging, quick culture medium exchange inside the incubator to reduce contamination risk and possibility to remove chips at different timepoints. In a second step, the set-up was used for running long-term (35 days) bone-remodeling-on-a-chip experiments.

Both osteoblasts and osteoclasts are essential to establish an *in vitro* bone remodeling model, as their interplay is important in the remodeling process [12]. As scaffold integration into a microfluidic device can be challenging [13], we relied on the cells’ own matrix production and mineralization. In the present study, human mesenchymal stromal cells (MSCs) were differentiated on-chip into osteoblasts that produced their own mineralized extracellular matrix and self-assembled into three-dimensional (3D) bone tissues. Next, human monocytes (MCs) were added and differentiated on-chip into osteoclasts, establishing a direct osteoblast-osteoclast coculture.

## 2. Materials and Methods

### 2.1 General experimental set-up

The general approach and timeline for the bone-remodeling-on-a-chip included several steps (Figure 1). Briefly, photolithography was employed as a fabrication method to create a silicon wafer mold, followed by soft lithography to create a polydimethylsiloxane (PDMS) layer in which the cell culture channels were located. The PDMS layer was bonded to a glass coverslip to complete the device. Before starting culturing cells in the device, the channels were coated with fibronectin to enhance cell adhesion. After the MSCs were well attached, osteogenic monoculture medium was perfused through the channels and culturing was continued for 21 days to allow for bone-like tissue formation. Next, MCs were added, and the medium was switched to coculture medium which was perfused through the channels for an additional 14 days.

**Figure 1.**
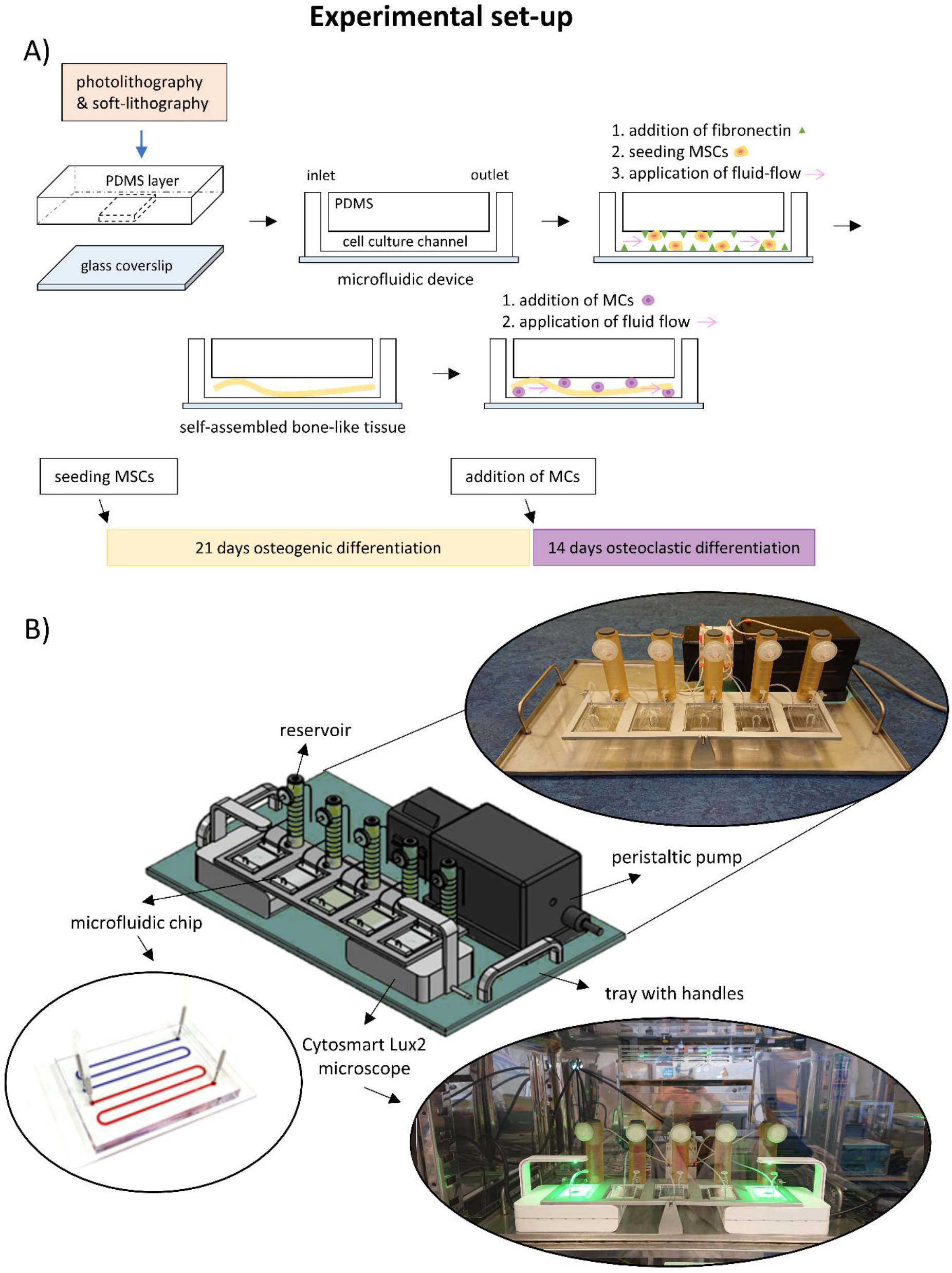
Outline of the experimental set-up. A) With the use of photo- and soft-lithography a PDMS microfluidic chip is created. The cell culture channel is coated with fibronectin 24 hours prior to seeding the MSCs. After 4 hours of attachment, culture medium fluid flow is applied for 21 days. The cells self-assemble into a bone-like tissue. After 21 days, MCs are seeded onto these tissues and allowed to attach for 24 hours. Next, fluid flow is applied, and the MCs are differentiated towards de osteoclastic lineage over an additional 14 days. B) The custom-made set-up comprises of a tray with handles on which a rack is fixed that holds five microfluidic chips and reservoirs. The peristaltic pump is placed behind this rack and Cytosmart Lux2 incubator microscopes are placed directly under two of the five chips.

### 2.2 Master mold fabrication and validation

A master mold for the patterned PDMS layer was produced with a silicon wafer by photolithography. The design of the photomask was drawn in AutoCAD (version 2021, Autodesk) and ordered at CAD/Art Services (Bandon, USA). First, a layer of negative photoresist (SU-8 2150, MicroChem, Newton, MA, USA) was spin-coated on top of a silicon wafer (Ø100 mm, Si-Mat). To ensure a thickness of 200 μm, the spin-coater (model WS-650MZ-23NPPB, Laurell, North Wales, USA) was set to a rotating speed of 2000 rpm for 30 seconds. The wafer was soft-baked, and the photomask was placed on top of the wafer, followed by UV-light (model UV-EXP150S-SYS, Idonus, Hauterive, Switzerland) exposure with a dose of 315 mJ/cm^2^, initiating SU-8 crosslinking of the exposed parts of the photoresist. A post exposure bake was done to complete the crosslinking process, followed by submerging the wafer into a developer solution (mr-Dev 600, Micro Resist Technology GmbH, Berlin, Germany) for 15 to 18 minutes to remove uncured photoresist. Subsequently, hard baking was performed to stabilize the printed pattern. The height of the channel dimensions was validated using a Mitutoyo Mu-Checker electronic comparator (model M402 519-402, Mitutoyo America Corporation, Aurora, USA). Finally, the master mold was silanized using 1H,1H,2H,2H-perfluorooctyltriethoxysilane (Fluorochem Ltd, Hadfield, UK) under vacuum overnight.

### 2.3 Microfluidic chip fabrication

The microfluidic chips consisted of a PDMS part and a PDMS-coated glass coverslip and were made by standard soft lithography. The PDMS part contained two separate meandering channels with each a dimension of 200 μm height × 800 μm width × 134 mm length. PDMS base and curing agent (Sylgard 184 silicone elastomer kit, Dow Corning, Midland, MI, USA) were thoroughly mixed at a 10:1 ratio (w/w), degassed under vacuum, and poured onto the wafer mold. After curing the PDMS overnight at 65°C, the patterned PDMS layer containing the channels was released from the master mold. A glass coverslip with a size of 35×64 mm and thickness of 0.17 mm was used to close off the microfluidic channels. To make sure the cells were exposed to the same substrate stiffness on the bottom surface as on the other walls of the channel, the coverslip was spin-coated with a thin layer of PDMS of approximately 130 μm in thickness. Rectangular PDMS pieces of 4 mm height × 35 mm width × 10 mm length were bonded at both ends of the PDMS chip to increase friction with the in- and outlet needles. Holes for the in- and outlets were punched with a Ø1.2 mm biopsy punch (Harris Uni-Core, No. 7093508). To attach the rectangular pieces to the patterned PDMS layer, 20 watt oxygen plasma was applied for 30 seconds using the plasma asher (model K1050X, Emitech, Quorum technologies, UK). Next, the coverslip was plasma bonded to the PDMS chip. The microfluidic device was completed by baking it in an oven at 65°C for 2 hours.

### 2.4 Design and fabrication of the perfusion set-up

The set-up was designed and fabricated in-house according to requirements that were set to enable long-term on chip cell culture (Figure 1B). These requirements included: easy transportation between safety cabinet and incubator, short tubing length between reservoir and chip, addition of live cell imaging devices, easy culture medium exchange and possibility to collect chips at different timepoints. The set-up was designed in Autodesk Inventor 2022 (detailed drawing in Supplementary Information Figure S1). The base plate was made of stainless steel and functioned as carry tray. The chip holder was made of aluminum and received anodizing surface treatment to make them more chemically resistant. The reservoirs were made of polysulfone and contained grooves that indicated the medium volume in mL. Each reservoir could hold up to 8 mL of liquid.

### 2.5 Assembly of the set-up and preparation of the microfluidic chips for cell culture

A closed perfusion-based set-up was constructed (Figure 1B). The set-up composed of a peristaltic pump (model P-70, Harvard Apparatus, Holliston, USA) and five in-house made polysulfone medium reservoirs. Each reservoir was connected to a sterile syringe filter (syringe filter 0.2 μm, CA, Sartorius) for air exchange and was closed off with a self-healing rubber injection port (Ø13 mm rubber bottle stoppers, SUCOHANS). Pump tubing (Ismatec pump tubing, 3-stop, PharMed BPT, 0.5 mm ID, yellow/orange) and silicon tubing (Ibidi GmbH, 0.5 mm ID) were connected to each other via metal tubes (Techcon, TE720050PK, 20G, 12.7 mm) and to the medium reservoirs via male luer connectors (Ibidi GmbH). Prior to assembling the set-up, the reservoirs and self-healing injection ports were sterilized with 70% ethanol and flushed with distilled water, and if possible, subsequently autoclaved. One day prior to cell seeding, the set-up was assembled, the reservoirs were filled with 6 mL sterile phosphate buffered saline (PBS). The pump was set at a speed of 100 μL/min to equilibrate the tubing and remove bubbles overnight. The set-up was placed in the incubator (37°C and 5% CO_2_). The microfluidic chips were sterilized with 70% ethanol and flushed 3 times with sterile PBS. Next, the chips were coated with fibronectin (Human Plasma Fibronectin Purified Protein, Merck, Schiphol-Rijk, The Netherlands) in sterile PBS (100 μg/mL) and placed overnight in the incubator (37°C and 5% CO_2_). At the day of cell seeding, the PBS was removed from the reservoirs, and they were filled with 6 mL cell culture medium each. The pump was set at a speed of 100 μL/min for at least 4 hours before connecting the cell seeded microfluidic chips to remove bubbles. The chips were flushed with sterile PBS three times and filled with cell culture medium prior to cell seeding.

### 2.6 Monoculture: isolation, expansion and cultivation of MSCs

MSC isolation and characterization from human bone marrow (Lonza, Walkersville, MD, USA) was performed as previously described [14]. MSCs were frozen at passage 3 with 1.25×10^6^ cells/ml in freezing medium containing fetal bovine serum (FBS BCBV7611, Sigma-Aldrich) with 10% dimethylsulfoxide (DMSO, 1.02952.1000, VWR, Radnor, PA, USA) and stored in liquid nitrogen until use. Before experiments, MSCs were thawed and seeded at a density of 2.5×10^3^ cells/cm^2^ in expansion medium containing DMEM (high glucose, 41966, Thermo Fisher Scientific), 10% FBS (BCBV7611, Sigma Aldrich), 1% Antibiotic Antimycotic (anti-anti, 15240, Thermo Fisher Scientific), 1% Non-Essential Amino Acids (11140, Thermo Fisher Scientific), and 1 ng/mL basic fibroblastic growth factor (bFGF, 100-18B, PeproTech, London, UK) at 37°C and 5% CO_2_. Upon 80% confluence, cells were detached using 0.25% trypsin-EDTA (25200, Thermo Fisher Scientific) and directly used for experiments at passage 4. Cells were resuspended at 1×10^6^ cells/mL in osteogenic monoculture medium containing DMEM (low glucose, 22320, Thermo Scientific), 10% human platelet lysate [15] (hPL, PE20612, PL BioScience, Aachen, Germany), 1% Anti-Anti, 0.1 μM dexamethasone (D4902, Sigma-Aldrich), 50 μg/mL ascorbic acid-2-phosphate (A8960, Sigma-Aldrich), 10 mM β-glycerophosphate (G9422, Sigma-Aldrich) and carefully pipetted into the channels of the chips. Each meandering channel contained ∼25 μL cell suspension and each chip had two meandering channels giving ∼50.000 cells per chip. Next, the seeded chips were incubated at 37°C and 5% CO_2_ for at least 4 hours before the pump was started to allow for cell attachment. Then the pump was set at a speed of 1 μL/min and the chips remained in the incubator for 21 days. Medium was refreshed weekly by removing 3 mL of the total 6 mL and replacing it by 3 mL fresh osteogenic medium with double the concentration of dexamethasone, ascorbic acid and β-glycerophosphate. Medium refreshment took place inside the incubator via the self-healing injection ports.

### 2.7 Coculture: isolation of MCs and cultivation of MSCs and MCs

Human peripheral blood buffy coats from healthy volunteers under informed consent were obtained from the local blood donation center (agreement NVT0320.03, Sanquin, Eindhoven, the Netherlands). The buffy coats (∼ 50 mL) were diluted to 200 mL in 0.6 % (w/v) sodium citrate in PBS adjusted to pH 7.2 at 4°C (citrate-PBS), after which the peripheral mononuclear cell fraction was isolated by carefully layering 25 mL diluted buffy coat onto 13 mL Lymphoprep (07851, StemCell technologies, Cologne, Germany) in separate 50 mL centrifugal tubes, and centrifuging for 20 min with lowest brake and acceleration at 800×g at room temperature. Human peripheral blood mononuclear cells (PBMCs) were collected, resuspended in citrate-PBS and washed 4 times in citrate-PBS supplemented with 0.01% bovine serum albumin (BSA, 10735086001, Sigma-Aldrich, Zwijndrecht, The Netherlands) to remove all Lymphoprep. PBMCs were frozen in freezing medium containing RPMI-1640 (RPMI, A10491, Thermo Fisher Scientific), 20% FBS (BCBV7611, Sigma-Aldrich) and 10% DMSO and stored in liquid nitrogen until further use. Prior to experiments, MCs were isolated from PBMCs using manual magnetic activated cell separation (MACS). PBMCs were thawed, collected in medium containing RPMI, 10% FBS (BCBV7611, Sigma-Aldrich) and 1% penicillin-streptomycin (p/s, 15070063, Thermo Fisher Scientific), and after centrifugation resuspended in isolation buffer (0.5% w/v BSA in 2mM EDTA-PBS). The Pan Monocyte Isolation Kit (130-096-537, Miltenyi Biotec, Leiden, The Netherlands) and LS columns (130-042-401, Miltenyi Biotec) were used according to the manufacturer’s protocol. After magnetic separation, the cells were directly resuspended in osteogenic coculture medium containing αMEM (41061, Thermo Fisher Scientific), 5% hPL, 1% Anti-Anti supplemented with 0.1 μM dexamethasone, 50 μg/mL ascorbic acid-2-phosphate, 10 mM β-glycerophosphate) spiked with 50 ng/mL macrophage colony stimulating factor (M-CSF, 300-25, PeproTech). After 21 days of culturing the MSCs, the MCs were added to establish the coculture. To seed the MCs into the chips, the entire set-up was placed in a safety cabinet and the tubing was carefully removed from the chips. The MCs were counted, and a suspension of 5×10^6^ cells /mL was carefully pipetted into the chips. Again, each meandering channel contained ∼25 μL cell suspension and each chip had two meandering channels giving ∼250.000 cells per chip. The monoculture medium was completely removed from the reservoirs and tubing and replaced with coculture medium. The chips were reconnected, and the setup was placed back into the incubator where cells were allowed to attach for 24 hours before the fluid flow was run again at 1 μL/min. After 2 days the coculture medium was replaced by coculture medium additionally containing 50 ng/mL receptor activator of NFκ-B ligand (RANKL, 310-01, PeproTech). Medium was changed weekly inside the incubator via the self-healing injection ports by removing 3 mL of the total 6 mL and replacing it by 3 mL fresh coculture medium with double the concentration of dexamethasone, ascorbic acid, β-glycerophosphate, M-CSF and RANKL. The culture was maintained for another 14 days, making a total of 35 days.

### 2.8 Brightfield time-lapse imaging

Brightfield images of the channels in the chip were taken to monitor cell morphology and assembly over time. One channel per chip and two chips per experiment were observed at 10× magnification with Lux2 microscopes (CytoSMART, Eindhoven, The Netherlands). Images were taken every three hours for the entire 35 days of culture.

### 2.9 Computational model for mechanical stimulation calculation

To quantify the mechanical stimulation in terms of fluid-induced wall shear stress and elastic strain on the cells, computational fluid dynamics (CFD) and fluid solid interaction (FSI) models were used (Figure 2). Based on experimental observation, the cells are flatly attached to the bottom of the microfluidic channel at day 0, while at day 21, the cells have detached from the bottom and have formed into a long 3D bone-like tissue. It was observed that the tissue occupied approximately 1/3 of the microfluidic channel width (i.e., 1/3 × 800 μm ≈ 267 μm), and the height was approximately 180 μm. In the computational model, the tissue geometry was idealized into an elliptical cylinder. As the tissue orientation could be either along or across the microfluidic channel at day 21, these two orientations were proposed (Figure 2B), and the tissue geometries were constructed in ANSYS DesignModeler (ANSYS Inc., Pennsylvania, USA). In the computational model, a representative volume of the channel (200 μm H × 800 μm W × 800 μm L) was modelled (Figure 2).

**Figure 2.**
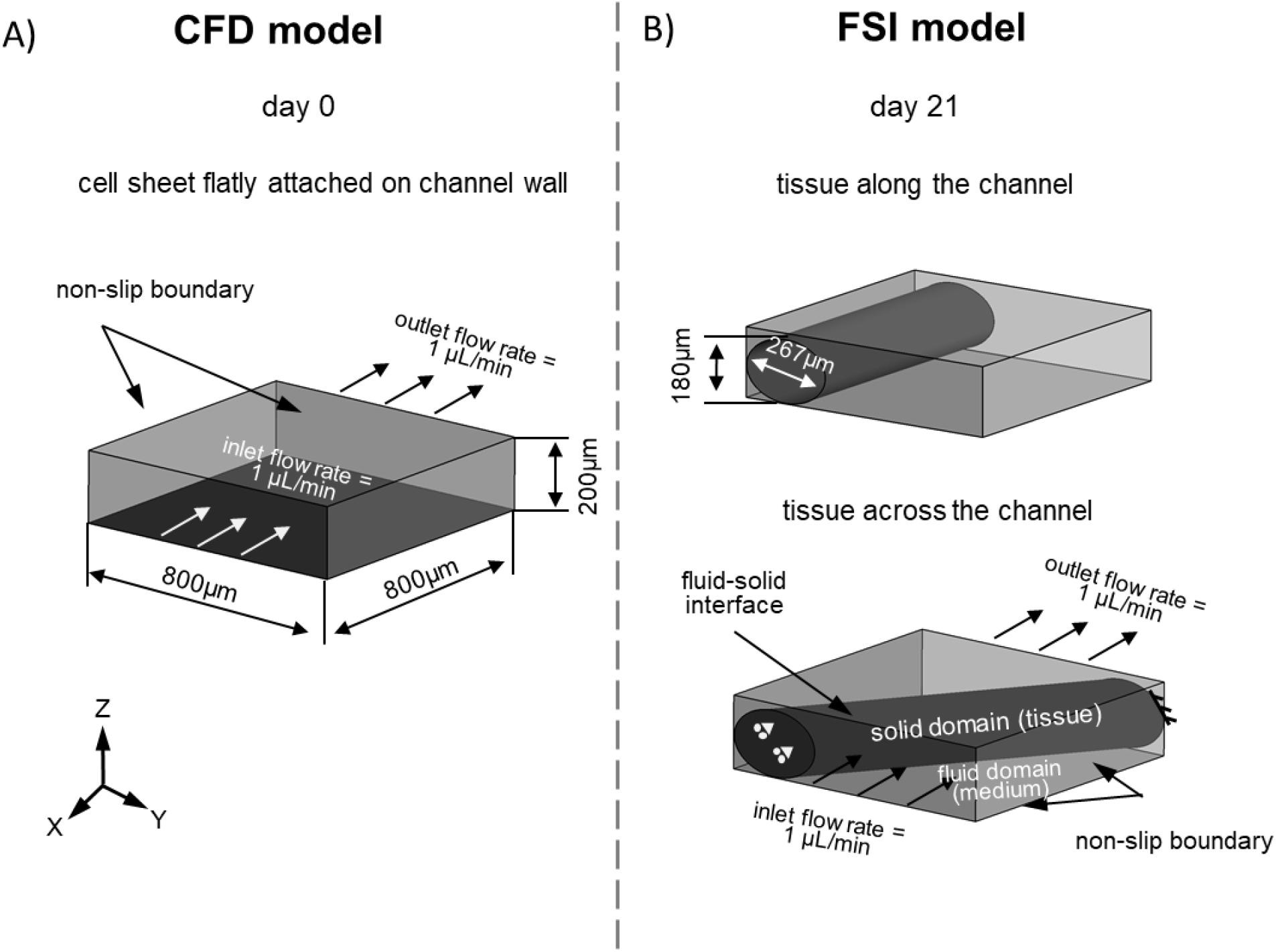
Determination of fluid shear stress. A) CFD model geometry and boundary conditions for calculating the shear stress on flat cell sheet at day 0. B) FSI model geometries and boundary conditions for calculating the shear stress on cells exposed to the medium and inside tissue at day 21.

For calculating the shear stress on cells at day 0, when the thin cell sheet was flatly attached on the channel bottom, the computational model was based on an empty channel by assuming that the shear stress on the channel bottom was equivalent to that on the cells. The calculation was based on the CFD model that follows the Navier-Stokes equation for incompressible flow:

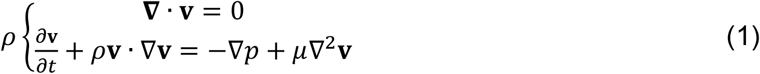

where, *ρ* and *μ* are medium density and dynamic viscosity, respectively (*μ* = 0.93 mPa·s, *ρ* = 1009 kg/m^3^ [16]); **v** is fluid velocity vector, *p* is pressure.

According to the pre-computation by ANSYS-CFX, the maximum Reynolds number was lower than 1. Therefore, the flow was defined as laminar flow. A flow rate of 1.0 μL/min used in experiment was prescribed at the inlet and outlet, respectively for mass conservation (Figure 2B). The channel walls were defined as non-slip boundaries (i.e., the fluid has zero velocity relative to the solid surfaces). The fluid domain was meshed by tetrahedral elements (global element size = 10 μm) with a patch conforming algorithm as described in Zhao *et al*. (2020) [17]. Moreover, the mesh for the cell/tissue region was refined with an element size of 4 μm, which generated 1,255,967 elements in total. The CFD model was solved using finite volume (FV) method by ANSYS CFX solver under the convergence criteria of root mean square (RMS) residual of momentum and mass < 1.0 × 10^−4^.

For calculating the fluid-induced wall shear stress on cells at the surface and internal strain of the bone-like tissue at day 21, a FSI model was used. The fluid domain was meshed with the same strategy as above, which generated 1,203,992 tetrahedral elements. The solid domain (bone-like tissue) was modelled as a hyperelastic (Neo-Hookean) material with the strain density function as equation (2), and solved by finite element (FE) method in ANSYS:

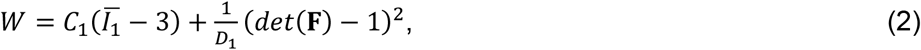

where *W* and **F** are the strain energy density and deformation gradient, respectively; 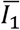 is the first invariant of the right Cauchy-Green deformation tensor, which is calculated as:

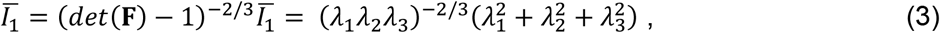

where *λ*_1_, *λ*_2_ and *λ*_3_ are the principal stretches.

In equation (2), *C*_1_ and *D*_1_ are material constants, which were calculated using shear modulus (*G*) and bulk modulus (*K*):

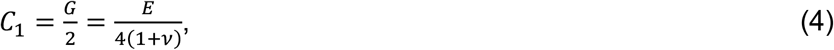

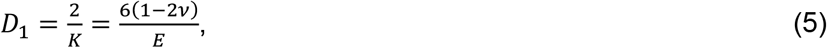

where *G, E* and *K* are shear modulus, Young’s modulus and bulk modulus, respectively; *v* is the Poisson’s ratio. The bone-like tissue (mixture of bone cells and extracellular matrix) was defined as an almost incompressible material with a Young’s modulus *E* of 1.70 kPa and Poisson’s ratio *v* of 0.49, according to the experimental measurements in Taiai *et al*. (2005) and Titushkin and Cho (2007) [18], [19]. So, the material parameters can be calculated as: *G* = 0.57 kPa, *K* = 28.33 kPa, *C*_1_= 0.29 kPa, *D*_1_= 7.00 × 10^−5^ Pa^−1^.

In terms of boundary conditions of the FE model, one end of tissue was fixed, and the other end was defined as frictionless support (Figure 2B). The tissue surfaces formed the fluid-solid interface between the CFD and FE domains and this two-way FSI analysis followed a staggered iteration approach, which coupled the fluid force and solid deformation as described in Zhao *et al*. (2020) [17].

### 2.10 Immunohistochemistry

At day 14, 21 and 35 chips were washed with PBS and fixed in 10% neutral-buffered formalin for 15 min. The chips were immunostained by washing with PBS-tween, permeabilizing in 0.5% Triton X-100 in PBS for 10 min and blocking in 10% normal goat serum in PBS for 30 min. Cells were incubated with DAPI, Phalloidin and immunostainings (Table 1) in PBS for 1 hour. Images were taken with either a fluorescent light microscope (Axio Observer 7, Zeiss, Oberkochen, Germany) (Figure 3B and 5A and B) or a confocal microscope (TCS SP5X, Leica Microsystems CMS GmbH, Mannheim, Germany) (Figure 5C).

**Table 1.** List of all dyes and antibodies used and their working concentrations/dilutions.

| Antigen | Source | Catalogue No | Label | Species | Concentration/Dilution |
| --- | --- | --- | --- | --- | --- |
| DAPI | Sigma-Aldrich | D9542 |  |  | 0.1 µg/mL |
| Atto 647 conjugated Phalloidin | Sigma-Aldrich | 65906 |  |  | 50 pmol |
| Collagen type-1 | Abcam | Ab34710 |  | Rabbit | 1:200 |
| RUNX-2 | Abcam | Ab23981 |  | Rabbit | 1:500 |
| Osteopontin | Thermo<br>Fisher | 14-9096-82 |  | Mouse | 1:200 |
| TRAP | Abcam | Ab185716 |  |  | 1:200 |
| Anti-rabbit<br>(H+L) | Molecular<br>Probes | A11008 | Alexa 488 | Goat | 1:200 |
| Anti-mouse<br>IgG1 (H+L) | Molecular<br>Probes | A21127 | Alexa 555 | Goat | 1:200 |

**Figure 3.**
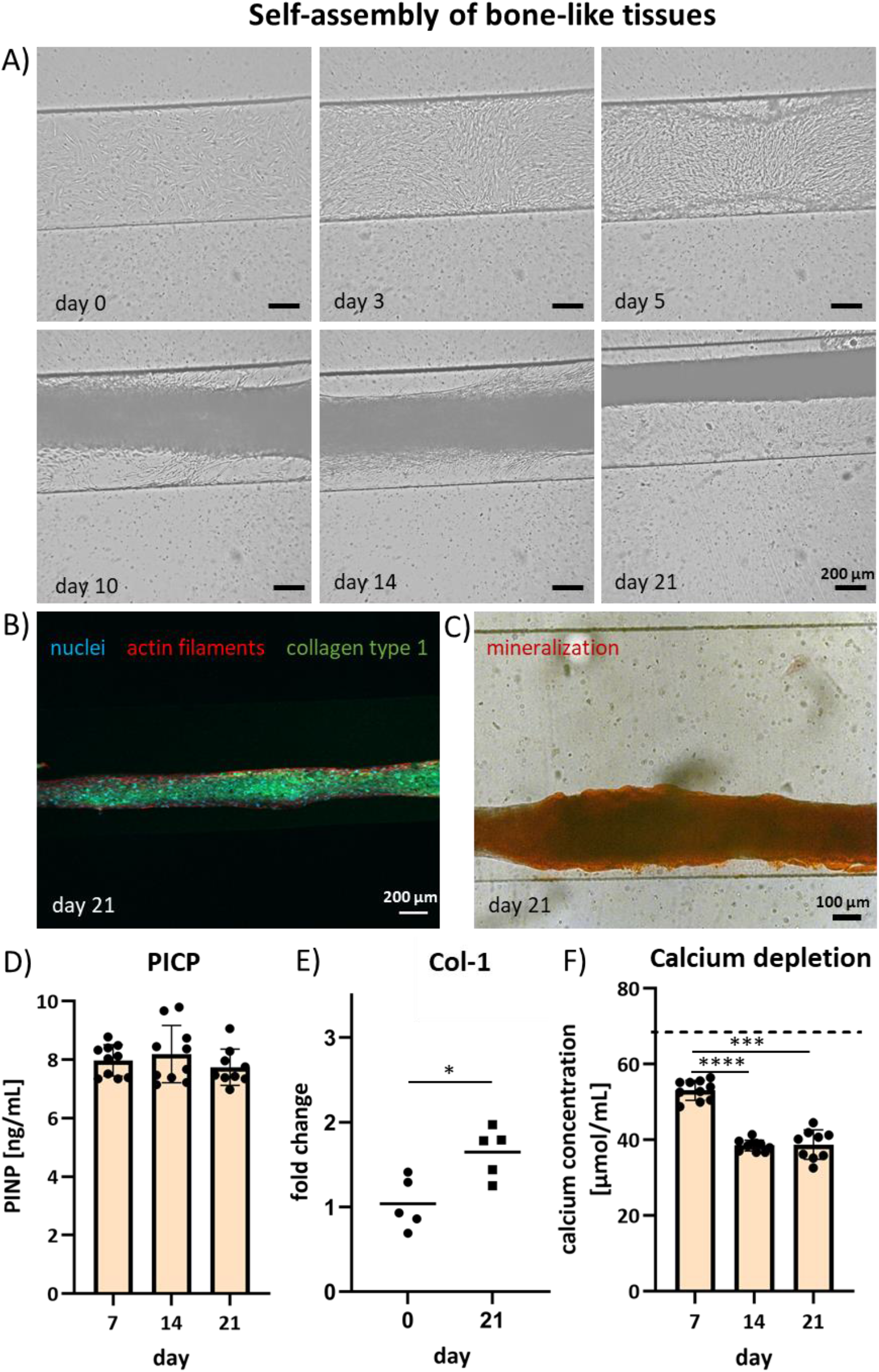
Self-assembly of 3D tissues. A) Time-lapse brightfield images of the 3D-self-assembly over time. At day 0, MSCs are seeded and attach to the channel in a monolayer. At day 3, the cells have spread over the bottom of the channel. At day 5, the cells start to detach from the channel walls. At day 10, the cells self-assemble into an elongated 3D tissue-like strut that becomes denser over the following days. B) The 3D tissue consists of collagen type 1 and C) is mineralized (Alizarin Red staining). D) PICP quantification as a measure for collagen formation (n=2 × 5). Differences are non-significant. E) Col-1 (n=5) gene expression measured before seeding in the chip (day 0) and after 21 days culturing in the chip (day 21). Values are displayed as 2^−ΔΔCt^ to day 0. Significant differences are shown by * for p<0.05. F) Calcium concentration in the culture medium showing calcium depletion (n=2 × 5). The dashed line represents the calcium concentration in fresh medium. Significant differences are shown by *** for p<0.001 and **** for p<0.0001.

**Figure 4.**
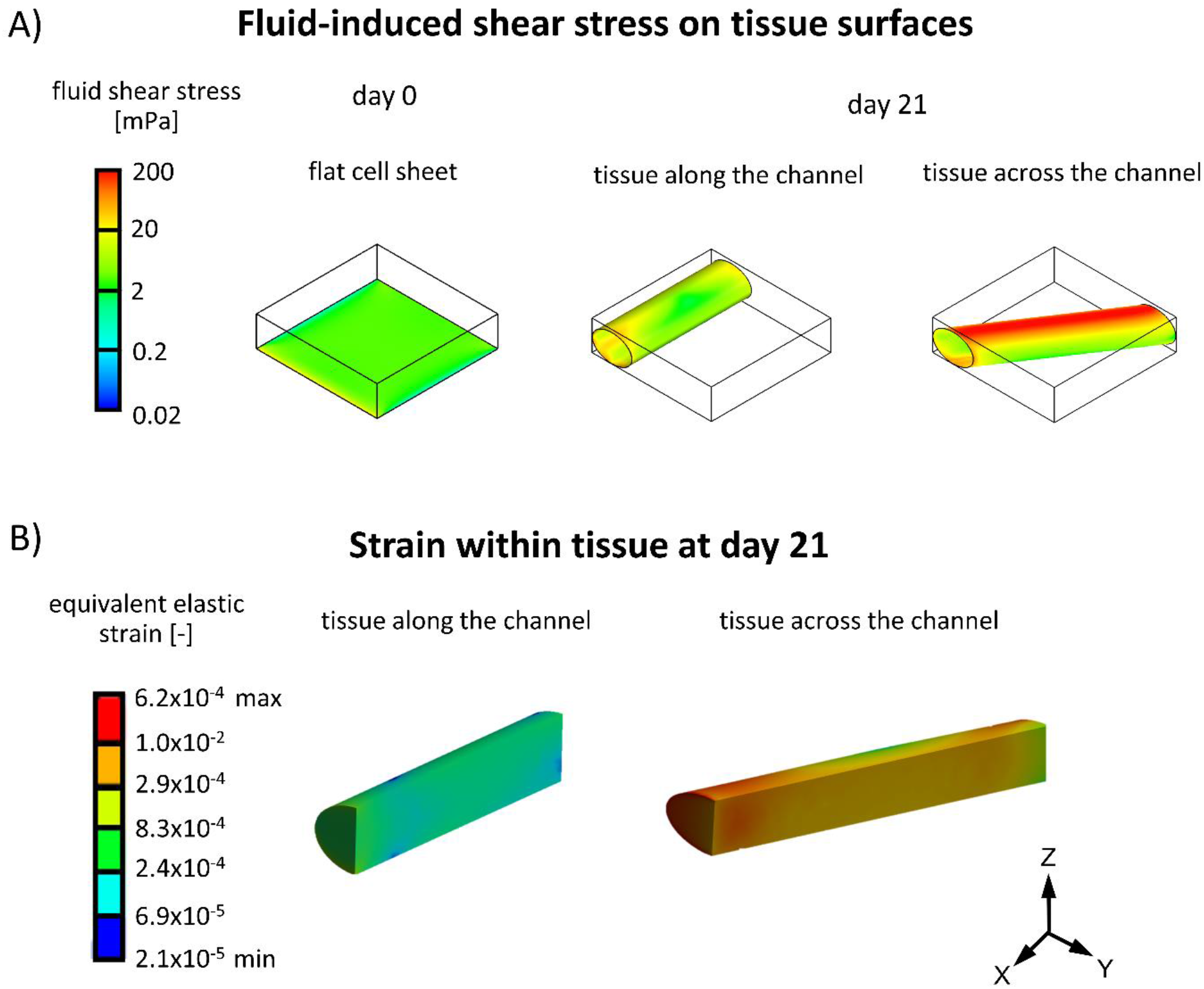
Shear stresses acting on the cells. A) Fluid-induced shear stress on cell/tissue surfaces at day 0 and 21. B) Sectional view of the shear stress within solid tissue (along and across the microfluidic channel) at day 21 due to the tissue deformation that is caused by fluid force.

**Figure 5.**
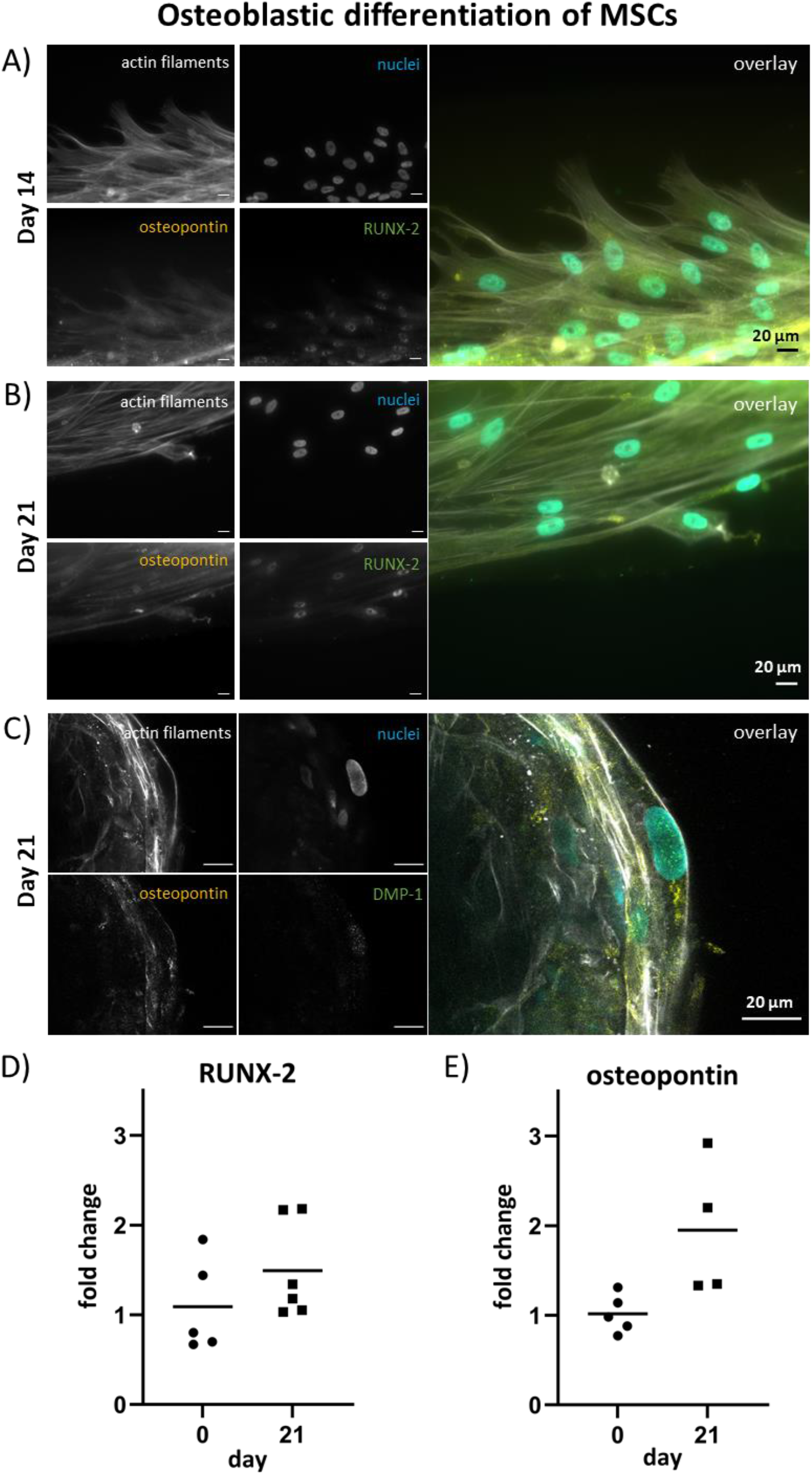
Osteogenic differentiation of MSCs: A) At day 14 and B) 21, osteopontin is expressed in the cells body and RUNX-2 in the nucleus. C) At day 21, DMP-1 expression is visible in the nucleus. All scale bars are 20 μm. D) RUNX-2 (n=5 or 6) and E) osteopontin (n=4 or 5) gene expression measured before seeding in the chip (day 0) and after 21 days culturing in the chip (day 21). Values are displayed as 2^−ΔΔCt^ to day 0. Differences are non-significant.

### 2.11 Histology

Following the immunostaining the same samples of day 14 and 21 were overstained with Alizarin Red (2% in distilled water, A5533, Sigma-Aldrich) for 15 minutes to identify mineralization. Subsequently, channels were washed with distilled water until no further discoloration of the water occurred. Images were made using a brightfield microscope (Axio Observer Z1, Zeiss, Oberkochen, Germany).

### 2.12 Quantification of Ca^2+^ concentration in supernatant

The calcium concentration from the supernatant was measured to determine changes in calcium concentration in the medium, as an indicator for mineralized matrix deposition/resorption during osteogenic differentiation. A calcium assay (Stanbio, 0150-250, Block Scientific, Bellport, NY, USA) was performed according to the manufacturer’s instructions (n=10). Briefly, 95 μl Cresolphthalein complexone reaction mixture was added to 5 μL sample and incubated at room temperature for 1 min. Absorbance was measured at 550 nm with a plate reader and absorbance values were converted to calcium concentrations using standard curve absorbance values.

### 2.13 Quantification of human pro-collagen 1 C-terminal propeptide

Human pro-collagen 1 C-terminal propeptide (PICP) as a product of collagen formation was quantified in cell supernatants of monocultured constructs at day 7, 14 and 21 using an enzyme-linked immunosorbent assay (ELISA, MBS2502579, MyBioSource, San Diego, CA, USA) according to the manufacturer’s protocol (n=2 × 5: two independent experiments with each 5 chips). 100 μL sample/standard was added to anti-human PICP coated microwells. After 90 min incubation at 37°C, samples were removed and replaced by 100 μL biotinylated antibody solution followed by 60 min incubation at 37°C. After thorough washing, 100 μL HRP-conjugate solution was added, and plates were incubated for 30 min at 37°C. Wells were again washed, and 90 μL substrate reagent was added followed by 15 min incubation in the dark at 37°C. To stop the reaction, 50 μL stop solution was added and absorbance was measured at 450 nm in a plate reader. Absorbance values were converted to PICP concentrations using standard curve absorbance values.

### 2.14 Gene expression by qPCR

To quantify gene expression levels, day 0 MSCs were pelleted by centrifuging 50.000 cells/pellet (230 rpm, 7 minutes), which is the same amount of cells as in the chips. Pellets were frozen and stored at -80°C. After 21 days of culturing in the chip, the self-assembled tissues were carefully taken out of the chips and also frozen and stored at -80°C. Frozen samples were crushed using a pestle to homogenize the samples, and subsequently lysed on ice using RLT lysis buffer. RNA was isolated using the Qiagen RNeasy kit (Qiagen, Hilden, Germany) following supplier instructions including a 15 minutes DNAse incubation step (Qiagen; 74106) to remove genomic DNA contamination. After extraction, RNA quantity and purity were assessed with a spectrophotometer (NanoDrop™ One, Isogen Life Science, The Netherlands). cDNA was synthesized in a thermal cycler (protocol: 65°C (5 min), on ice (2 min) while adding the enzyme mixture, 37°C (2 min), 25°C (10 min), 37°C (50 min), and 70°C (15 min)) starting from a 20 μL reaction solution containing 200 ng of RNA, 1 μL dNTPs (10 mM, Invitrogen), 1 μL random primers (50 ng/μL, Promega, C1181), 2 μL 0.1 M DTT, 4 μL 5× first strand buffer, 1 μL M-MLV Reverse Transcriptase (RT) (200 U/μL, Invitrogen, 28025-013, Breda, the Netherlands) and supplemented with RNAse-free ultra-pure water (ddH2O). Genomic DNA contamination was checked with glyceraldehydes-3-phosphate dehydrogenase (GAPDH) primers, conventional PCR, and gel electrophoresis.

qPCR was executed to investigate the expression of genes related to osteogenic differentiation: COL-1, RUNX-2 and SPP1 (osteopontin), utilizing the primer sequences listed in Supplementary Information Table S1. Six reference genes were tested of which the two most stable ones were used (ATP5F1B and TOP1). Expression was investigated by adding 500 nM primer mix, 5 μL SYBR Green Supermix (Bio-Rad; 170-8886), and an additional 1.75 μL ddH2O to 3 μL of diluted cDNA. C_t_ values were acquired by exposing the mixtures to the following thermal protocol: 95°C (3 min), 40 cycles of 95°C (20 s), 60°C (20 s), and 72°C (30 s), 95°C (1 min), and 65°C (1 min), concluded with a melting curve measurement. Differences in expression profiles were determined by normalizing the C_t_ values for the reference gene (only ATP5F1B shown, similar results to TOP1) (ΔCt), correcting these values for the C_t_ value of the control (day 0) (ΔΔC_t_) and applying the 2^−ΔΔCt^ formula to determine the fold changes in expression.

### 2.15 Quantification of tartrate-resistant acid phosphatase activity in the supernatant

Tartrate-resistant acid phosphatase (TRAP) concentration from the supernatant of the coculture was measured at day 23, 28 and 35 to determine the amount of secreted TRAP during osteoclastic differentiation (n=2+3: two independent experiments, one with 2 and one with 3 chips). 10 μL of supernatant was placed in a 96-well plate and resuspended in 90 μL assay buffer containing 3M NaAc, 10% Triton X-100, 1 mg/mL p-nitrophenyl phosphate (pNPP, Sigma Aldrich, 71768) (pH=5.5). Samples were incubated for 1.5 hours at 37°C. Finally, 100 μL 0.3M NaOH solution was added to stop the reaction. The amount of TRAP was determined via the optical absorbance at 405 nm and absorbance values were converted to TRAP activity using standard curve absorbance values.

### 2.16 Quantification of human crosslinked C-telopeptide of collagen type 1

Human crosslinked C-telopeptide of collagen type 1 (CTX) as a collagen degradation product was quantified in cell supernatants of cocultured constructs at day 23, 28 and 35 using an ELISA (MBS162789, MyBioSource, San Diego, CA, USA) according to the manufacturer’s protocol (n=2+3: two independent experiments, one with 2 and one with 3 chips). 50 μL standard and 40 μL sample were separately added to anti-human CTX coated microwells. 10 μL Anti-CTX antibody was added to each sample well and 50 μL streptavidin-HRP was added to all sample and standard wells and incubated for 60 min at 37°C. After thorough washing, 50 μL substrate reagent A and 50 μL substrate reagent B were added, followed by 10 min incubation in the dark at 37°C. To stop the reaction, 50 μL stop solution was added and absorbance was measured at 450 nm in a plate reader. Absorbance values were converted to CTX concentrations using standard curve absorbance values.

### 2.17 Statistical analysis

Statistical analyses were performed, and graphs were prepared in GraphPad Prism (version 9.4.0, GraphPad, La Jolla, CA, USA) and R (version 4.0.2). Data was tested for normal distribution with Shapiro-Wilk tests. All data was normally distributed. One-way repeated measures ANOVA was performed for PICP, calcium, TRAP and CTX to compare consecutive timepoints. Post-hoc Tukey was applied to correct for multiple comparisons. qPCR data was tested for equal variances using Levene’s test. As variances were not equal, Welch’s t-test was performed for qPCR data to compare day 0 to day 21. All data is presented as mean plus/minus standard deviation.

## 3. Results

### 3.1 Experimental set-up enables long-term on-chip cell culture

A set-up was designed that successfully facilitated long-term (35 days) on-chip cell culture (Figure 1B). The set-up was quick to assemble. The carry tray with handles provided easy and safe transportation between safety cabinet and incubator. All components had a fixed position, refraining movement during transportation. The reservoir and chip were placed approximately at same height to reduce pressure difference, which has been described to prevent a distorted flow [7]. Inside the incubator, the reservoirs were easily accessible for safe and fast culture medium exchange through the self-healing rubber injection ports. The short tubing between reservoir and chip (2.5 cm) reduced the risk of air bubble formation, as long tubing length increases the assembly of small air bubbles into large ones. Compared to our previous set-up we were able to reduce the total tubing length by 46%, decreasing the bubble formation risk. Consequently, double the amount of cell seeded chips survived the 35-day culture period (detailed comparison with our previous set-up in Supplementary Information Table S2 and Figure S2).

### 3.2 Self-assembly of elongated 3D tissues containing of collagen type 1 and minerals

The morphology and behavior of cells was investigated using time-lapse live cell imaging (Figure 3A). The bright-field images showed that at day 0, the MSCs were spindle-shaped, which confirmed their attachment to the channel surfaces. At day 5, cells started to detach from the channel side walls. At day 10, cells self-assembled, thereby forming a string of connected cells in the middle part of the channel. The string compacted over time. At day 21, a dense 3D elongated construct was visible. Over the entire culture period, some cells remained attached to the channel walls to keep the formed tissue in place. This proved to be essential to withstand the shear stress created by the fluid flow. Non-attached cells and cell-assemblies were flushed out of the channels. At day 21, the self-assembled constructs were investigated for their composition. The extracellular matrix produced by the cells consisted of collagen type 1 (Figure 3B) and was highly mineralized as shown by Alizarin red staining (Figure 3C). The rate of collagen type 1 formation stayed constant over time shown by measuring PICP in the culture medium (Figure 3D). Collagen type 1 gene expression was significantly higher at the day 21 osteoblasts compared to day 0 MSCs (Figure 3E).

Quantification of calcium concentration in the cell culture supernatant showed that calcium was depleted from the supernatant during the first 21 days (Figure 3F). Significantly more calcium was depleted on day 14 and 21 compared to day 7. This observation together with the histological stainings implies that the produced collagenous extracellular matrix was mineralized with a calcium mineral and indicates the formation of a self-assembled 3D bone-like tissue.

### 3.3 Mechanical stimulation increased upon self-assembly of the tissues and is dependent on tissue orientation

Computational models (CFD and FSI) were used to quantify the mechanical stimulation in terms of shear stress and mechanical strain received by the cells in the chip. The fluid-induced shear stress on the monolayer of cells at day 0 was 3.66 mPa (Figure 4A). At day 21, the self-assembled cells on the surface received an average shear stress of 10.55 mPa and 45.30 mPa respectively for the tissue oriented along and across the channel (Figure 4A). Due to the deformation of the tissue under fluid flow, the cells embedded within the tissue could experience the mechanical strain. The average values of equivalent elastic strain within the tissue were 4.74 × 10^−4^ and 6.65 × 10^−3^ respectively for the tissue oriented along and across the channel (Figure 4B). These results show that the fluid flow shear stress on the cells both on the surface and within the tissue increased upon self-assembly and is highly dependent on the orientation of the self-assembled tissue.

### 3.4 Osteogenic differentiation of MSCs in a 21 day monoculture

To confirm osteoblastic differentiation of the MSCs in the first 21 days, samples were stained for RUNX-2 and osteopontin at day 14 and 21 and for DMP-1 at day 21. RUNX-2 was present in the nucleus and osteopontin in the body of the cells at day 14 and 21 (Figure 5A and B). DMP-1 was visible in the nucleus of the cells at day 21, indicating early signs of transition to osteocytes (Figure 5C). RUNX-2 and osteopontin gene expression demonstrated an upward trend comparing day 21 to day 0 (Figure 5D and E). The results confirm that the microfluidic chip allows for MSCs to differentiate into osteoblasts over time. It is to be noted that the density of the self-assembled 3D bone-like tissue complicated the imaging process, as the laser light was not able to penetrate through the thick matrix. Only cells on the surface of the tissues could be visualized.

### 3.5 MCs fuse into multi-nucleated cells in 14-day coculture

After the 21-day tissue formation phase, MCs were added to the microfluidic chips and cultured for another 14 days (35 days in total). Monoculture medium was changed into coculture medium containing RANKL and M-CSF to induce osteoclastic differentiation of the MCs. At day 35, Z-stack images revealed the presence of mono- and multi-nucleated cells with different morphologies (Figure 6A and B). Elongated mononuclear cells with large (± Ø 16 μm) oval shaped nuclei were identified as osteoblasts (Figure 6A, orange arrow). Round, multinucleated cells with smaller (± Ø 12 μm) round nuclei were identified as fused monocytes/pre-osteoclasts (Figure 6A, purple arrow). An additional TRAP immunostaining revealed a multi-nucleated cell with a clear actin-ring and expression of TRAP in the cell body, found on the surface of the 3D bone-like tissue (Figure 6B). These results suggest that MCs were able to attach to the tissues upon cell seeding and could withstand the application of fluid flow within the chip. The MCs were stimulated towards the osteoclastic lineage shown by the formed multi-nucleated cells and TRAP staining.

**Figure 6.**
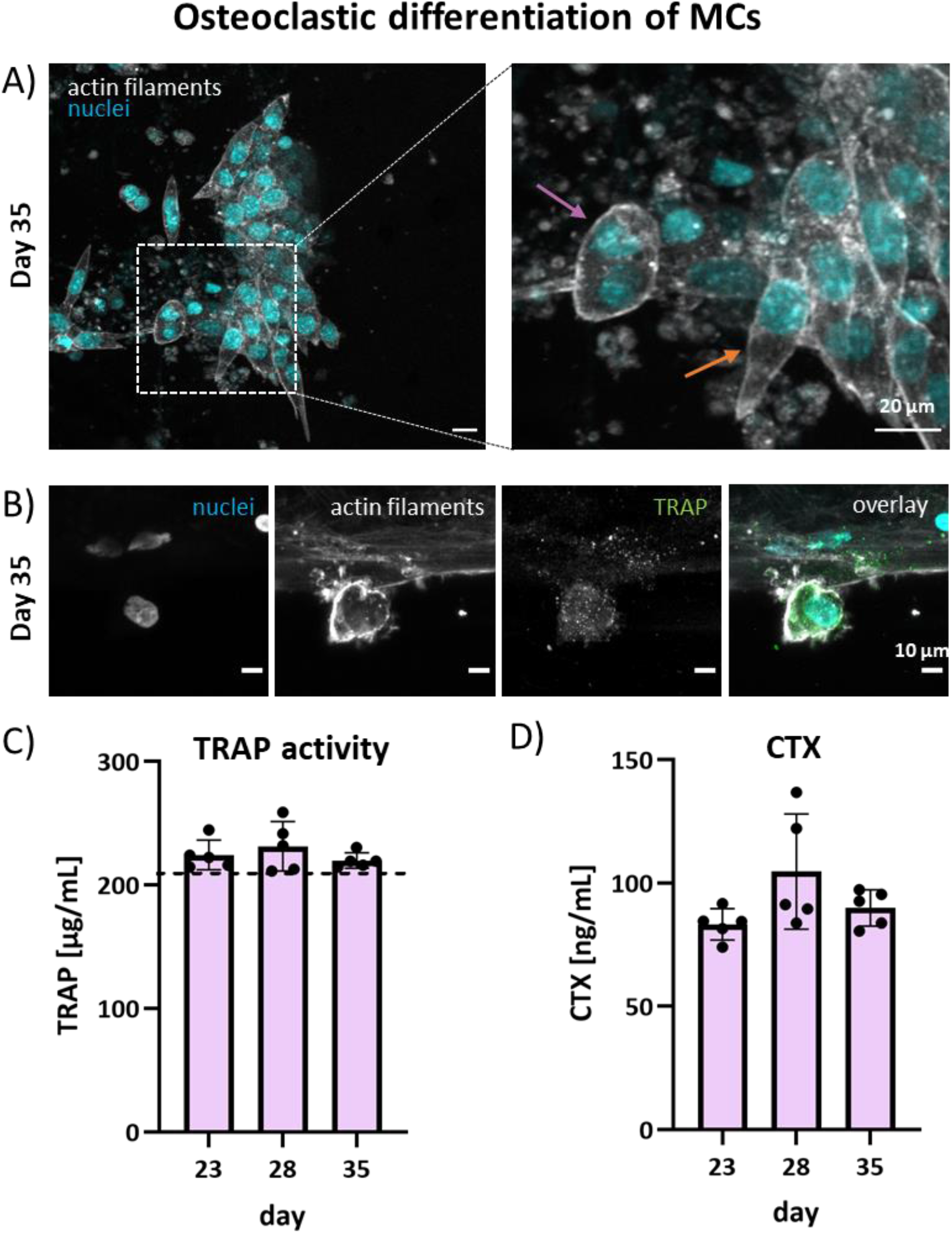
Osteoclastic differentiation of MCs within the coculture. A) At day 35, round multinucleated cells (purple arrow) and elongated mononuclear cells (orange arrow) are visible. B) On the surface of the bone-like tissue, a round, multinucleated cell is visible with a clear actin ring and TRAP expression in the cells body. This image is a maximum projection of z-stacks. C) TRAP activity (n=2+3); Dashed line represents the TRAP activity in fresh medium, containing TRAP originating from the hPL. Differences are non-significant. D) CTX quantification as a measure for collagen degradation (n=2+3). Differences are non-significant.

TRAP activity was measured in the coculture medium over time (Figure 6C). Differences between timepoints were non-significant. However, the TRAP activity measured in the samples showed a trend of being higher than the activity measured in fresh culture medium (Figure 6C, dashed line), indicating that there might be some contribution from the cells. The TRAP activity in fresh medium is most likely originating from the hPL [15].

By measuring CTX in the medium, collagen degradation by the cells was quantified. Collagen type 1 degradation was similar over time with non-significant differences between the timepoints (Figure 6D). A slight trend towards higher degradation at day 28 was seen.

## 4. Discussion

A bone-on-a-chip coculture system was developed in which human MSCs differentiated into osteoblasts and self-assembled into scaffold-free bone-like tissues with the shape and dimensions of human trabeculae (100–200 μm thick cylindrical rods) [33]. Human MCs were able to attach to these tissues and to fuse into multinucleated osteoclast-like cells, establishing the coculture. The microfluidic chip and set-up were able to maintain the culture over a 35 day culture period. This on-chip coculture is a first step towards *in vitro* bone remodeling models for drug testing.

To date, bone-on-a-chip models have not been abundantly reported, probably because of the highly complex bone microenvironment and multicellularity [6]. Coculturing osteoblasts and osteoclasts within the desired timeline is difficult given the differing differentiation timelines and cell culture medium compositions needed [9], [20]. Nonetheless, bone-on-a-chip systems have been reported for example for bone cell signaling [21], mechanical stimulation [22] and diseases [23], [24]. Most studies use cell lines and/or animal cells and do not incorporate a direct coculture of the different bone cells. Our specific *in vitro* bone remodeling model is fully human, having the advantage of no interspecies differences [25], [26], [27], [28]. We used primary cells, avoiding cell lines which are manipulated to enable continuous passaging, possibly affecting their outcomes [29].

To our knowledge, only one other research group has reported a bone-on-a-chip design specifically for bone remodeling [29]–[32]. Their lab-on-a-chip device comprises of three wells with each one of the three bone cell types. The osteoblasts (MC3T3-E1) are seeded on polystyrene discs, the osteoclasts (RAW 3264.7) are seeded on bone wafers, and osteocytes (MLO-Y4) on collagen type 1. The channels between the wells allow for exchange of conditioned medium. The chip allows for mechanical loading of the osteocytes by applying a static out of plane distention to stretch the cells [31], [32]. This platform allows for many different configurations and is planned to be used with direct cocultures and fluid flow. To date only indirect coculture and cell lines are used. While indirect cocultures allow cell-cell communication though soluble factors, the communication through their surface receptors and gap junctions is missing [20]. We show a direct coculture using primary cells and mechanical stimulation. Direct communication between osteoblasts and osteoclasts can be beneficial for the bone remodeling process as for example next to the soluble factors M-CSF and RANKL also their membrane bound variants play an important role in osteoclastogenesis [12], [33].

Mechanical loading is a critical environmental factor during bone development and homeostasis. *In vivo*, bone shape, mass, and trabecular architecture change constantly in response to mechanical loading, a process called bone adaptation [34]. Shear stress and mechanical strain are both examples of mechanical loading that occurs in bone. Many types and magnitudes of shear stress and mechanical strain have been shown to have positive effects on bone cells. For example, fluid flow shear stress has been show to enhance the osteogenic differentiation of MSCs [34]. A broad range of fluid flow shear stresses in microfluidic devices for bone tissue engineering has been reported from 0.01 mPa to 1.03 Pa [35]–[38]. Our system falls into this range with fluid flow shear stresses of 3.66 mPa up to 45.30 mPa (Figure 4). The cells on the surface of the self-assembled 3D tissue are expected to feel shear stresses up to 45.30 mPa (Figure 4A). These cells are most likely the progenitor cells, osteoblasts, bone lining cells and osteoclasts. The embedded cells, the osteocytes, are expected to feel elastic strain up to 6.65 × 10^−3^ (Figure 4B). For comparison, during walking bones experience strains around 1 × 10^−4^ and *in vivo*, strain in the range of 1 × 10^−4^ – 2 × 10^−3^ is shown to be optimal for bone healing [39]. It needs to be noted that our model used for the embedded cells, is based on a number of assumptions and did for example not take into account the hardening of the tissue when it mineralizes, which in turn might affect the load on the embedded cells. In future experiments it would be interesting to take a closer look to both the influence of tissue orientation (and thus altered mechanical stimulation) on cell differentiation and matrix production, and to the effects of mineralization on strain.

The analysis methods on chips and thus also in our chip can be challenging. At the moment, most methods are off-chip endpoint measurements, while non-destructive methods would be desired [6], [12]. Our system is still restricted in the number of samples we can generate, and the extended culture time reduces our ability to produce the desired number of samples necessary to carry out a complete analysis. Our set-up does offer the advantage of medium sampling thanks to the self-healing injection ports. In this way, assays on the culture medium can be performed at different timepoints. However, the dilution of secreted growth factors in a microfluidic chip using large media reservoirs in combination with the small amount of cells present in the chip is also a concern as it asks for sensitive assays [40]. In the future, integration of biosensors into the chip could provide sensitive, non-invasive continuous monitoring of the experiment [6]. For example, efforts in developing microfluidic based biosensing technologies for continuous and long-term measurement of glucose, lactate, pyruvate, dissolved oxygen, pH and reactive oxygen species have already been reported [41].

In bone, the extracellular matrix produced by the cells is a very important microenvironment that is particularly essential for the differentiation of osteoblasts into osteocytes. To resemble physiological bone, the organic matrix should comprise a highly dense and aligned collagen network [42]. Mineralization should occur inside and outside of the collagen fibrils [43]. In our system, collagen appears to orient in the fluid-flow direction (Figure 3B) and mineralization of the extracellular matrix was observed (Figure 3C). However, to ensure proper collagen alignment and correct mineral location, advanced techniques such as transmission electron microscopy (TEM) and focused ion beam scanning electron microscopy (FIB-SEM) that allow for nanoscale sample investigation are needed [42]. Future work is needed to elucidate the nanoscale properties of our matrix.

When the challenges of sample size and analytical methods have been overcome in the future, our bone-remodeling-on-a-chip is expected to provide valuable platform for drug testing. In comparison to cell monocultures, direct coculture offers the opportunity to study the interaction between osteocytes, osteoblasts and osteoclasts under influence of different stimuli, such as drugs and altered mechanical loads in a 3D environment resembling bone trabeculae. We envision the potential to use both healthy donor cells and patient cells in the model to compare the effects of drugs. Furthermore, we would be able to take steps towards personalized medicine and accurately screen for the most suitable drugs for individual patients.

In conclusion, we have demonstrated a scaffold-free, fully based on primary human cells, 3D microfluidic coculture model. MSCs were differentiated on-chip into osteoblasts and self-assembled into bone-like tissues with the dimensions of human trabeculae. Next, MCs were added to these tissues and differentiated on-chip into osteoclast-like cells. Furthermore, a set-up was developed allowing for long-term (35 days) on-chip cell culture with benefits including mechanical stimulation through applied fluid-flow, low bubble formation risk, easy culture medium change inside the incubator and live-cell imaging options. This on-chip coculture is a crucial advance towards developing *in vitro* bone remodeling models to facilitate drug testing in the future.

## 7. Author contributions

MV, KI and SH contributed to conception, methodology and design of the study. MV, EB and MB performed the experiments. MV analyzed the results. FZ made and analyzed the computational model. JB, HEA, MV and MB designed the set-up. JB drew and built the set-up. MvD executed the RNA isolation, cDNA synthesis and qPCR and analyzed the gene expression results. MV wrote the original draft of the manuscript and prepared the figures. All authors contributed to manuscript revision and approved the submitted version. KI and SH contributed to the supervision. SH acquired funding for this research.

## 8. Acknowledgements

We thank dr. Bregje de Wildt for her expertise and help on the coculture experiments. We thank dr. Andreas Pollet for his expertise and help on the microfluidic cell culture.

