## Supplementary Information for "Towards bone-remodeling-on-a-chip: self-assembling 3D osteoblast-osteoclast coculture in a microfluidic chip"

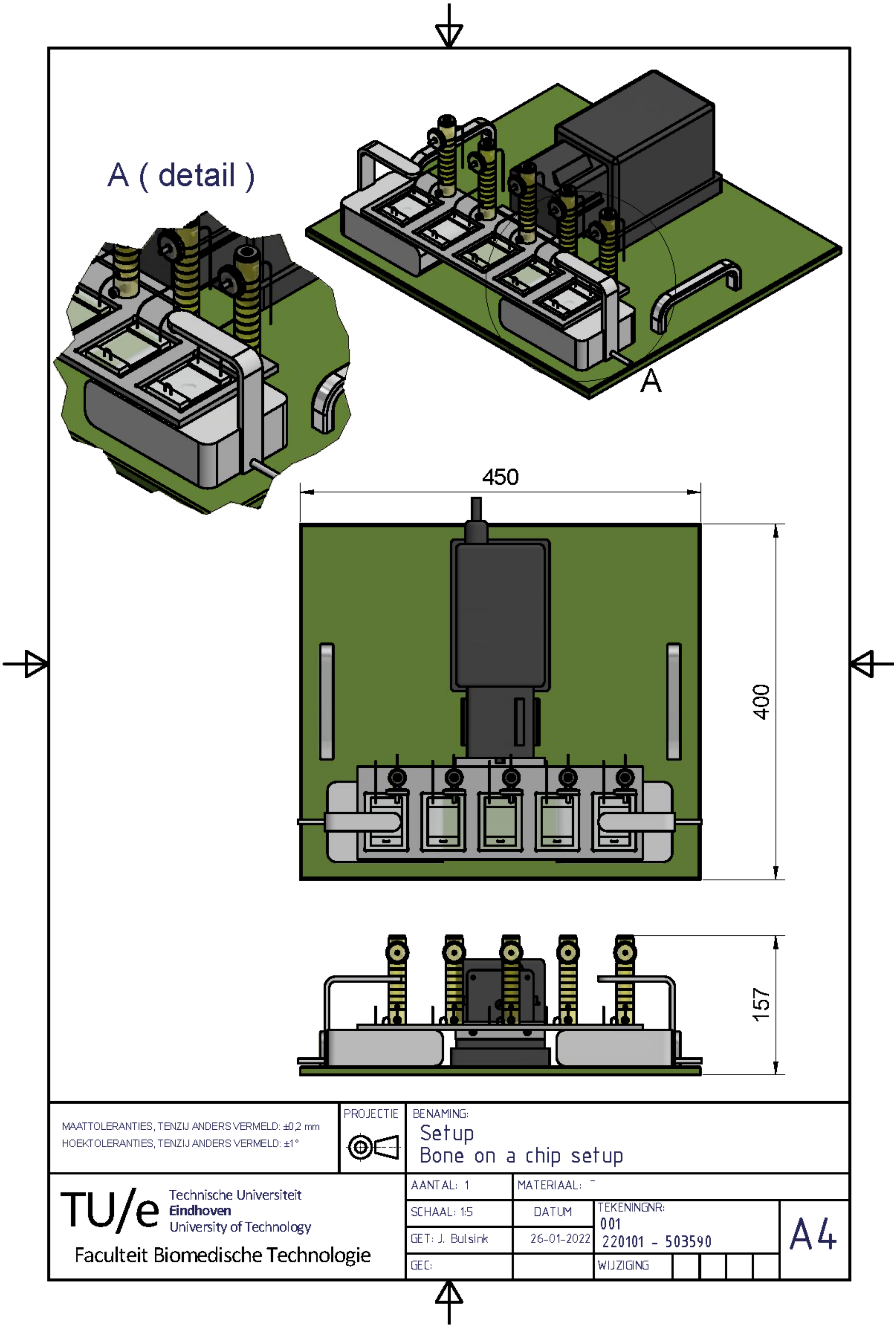


*Figure S1. Detailed drawing of the optimized set-up. Measures are in millimeter.*

*Table S1. Primer sequences of analyzed genes*

| Gene | 5' – 3' |
|---|---|
| COL1 | F- AATCACCTGCGTACAGAACGG<br>R- TCGTCACAGATCACGTCATCG |
| RUNX2 | F- GTCATGGCGGGTAACGATGA<br>R- GGGTTCCCGAGGTCCATCTA |
| SPP1 | F- GCCGAGGTGATAGTGTGGTT<br>R- AACGGGGATGGCCTTGTATG |
| ATP5F1B | F- CCAGCAGATTTTGGCAGGTGA<br>R- AGACCCCTCACGATGAATGC |

*Table S2. Comparison of performance of initial and optimized set-ups.*

| Measured parameter | Initial set-up | Optimized set-up |
|---|---|---|
| Duration of assembling the setup inside biosafety cabinet | ~1.5 - 2h | ~45 min - 1h |
| Total tubing length | 51 cm | 23.5 cm |
| Tubing length between reservoir and chip | 15 cm | 2.5 cm |
| Number of connectors and valves | 6 | 5 |
| Duration of medium change | 30 min | 15 min |
| Chip cell survival after culture period | ~44% | ~90% |

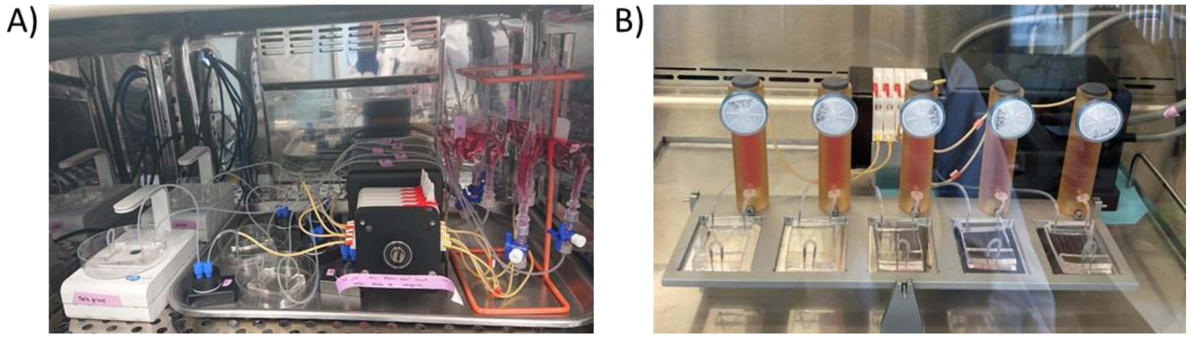


*Figure S2. Photographic images of A) the initial set-up and B) the optimized set-up showing the difference in level of organization, tubing length and handleability.*
